# Protein kinase PfPK2 mediated signalling is critical for host erythrocyte invasion by malaria parasite

**DOI:** 10.1101/2023.07.31.551280

**Authors:** Rahul Singh Rawat, Ankit Gupta, Neelam Antil, Sonika Bhatnagar, T.S. Keshava Prasad, Pushkar Sharma

## Abstract

Signalling pathways in malaria parasite remain poorly defined and major reason for this is the lack of understanding of the function of majority of parasite protein kinases and phosphatases in parasite signalling and its biology. In the present study, we have elucidated the function of Protein Kinase 2 (PfPK2), which is known to be indispensable for the survival of human malaria parasite *Plasmodium falciparum*. We demonstrate that it is involved in the invasion of host erythrocytes, which is critical for establishing infection. In addition, PfPK2 may also be involved in the maturation of the parasite post-invasion. PfPK2 regulates the release of microneme proteins like Apical Membrane Antigen 1 (AMA1), which facilitates the formation of Tight Junction between the merozoite and host erythrocyte-a key step in the process of invasion. Comparative phosphoproteomics studies revealed that PfPK2 may be involved in regulation of several key proteins involved in invasion and signalling. Furthermore, PfPK2 regulates the generation of cGMP and the release of calcium in the parasite, which are key second messengers for the process of invasion. These and other studies have shed light on a novel signalling pathway in which PfPK2 acts as an upstream regulator of important cGMP-calcium signalling, which is plays an important role in parasite invasion.

## Introduction

Malaria contributes to almost 400,000 deaths globally (Hay et al, 2004). While there is a significant reduction in both number of malaria cases and deaths, the efforts to eradicate this disease have been threatened by the emergence of resistance exhibited by malaria parasite-the most lethal species *Plasmodium falciparum*- against commonly used drugs (Mathenge et al, 2020). Therefore, new therapeutic inventions are needed to keep malaria under control and prevent its rise again. It is the asexual development of the parasite in blood stages that results in malaria pathology (Miller et al, 2002). The blood stage development of the parasite starts with the invasion of host erythrocyte by the merozoites. Subsequent to its entry the parasite forms a parasitophorous vacuole (PV) within which parasite develops from early ring stages to trophozoites, replicates its genome and undergoes asynchronous asexual division in schizont stages. Following segmentation, individual merozoites are generated, which egress after rupture of the host erythrocytes and freshly released merozoites invade new erythrocytes. Erythrocyte invasion is a key step in establishing the infection, therefore, an important process to target for which deep insights into molecular mechanisms are needed (Cowman & Crabb, 2006). The merozoite has highly specialized organelles- micronemes, rhoptries and dense granules- at its apical end that secrete proteins that serve as ligands for receptors present at the erythrocyte surface. Several receptor-ligand interactions are involved in facilitating the process of invasion, which is a tightly regulated process (Cowman et al, 2017). After initial contact and attachment, the merozoite reorients to position its apical end for rhoptries to secrete their material. In addition to interacting with receptors on host surface, some microneme proteins like Apical Membrane Antigen-1 (AMA1) are also involved in the formation of Tight Junction with erythrocyte. A family of rhoptry neck proteins (RONs) are also critical for merozoite entry into the erythrocyte (Lamarque et al, 2011; Srinivasan et al, 2011) as some of the RONs interact with microneme secreted AMA1 present on the merozoite surface, which is critical for the formation of Moving or Tight Junction (TJ) between the parasite and the host erythrocyte (Richard et al; Srinivasan et al, 2011). The actomyosin motor is postulated to provide the force necessary to move the TJ formation towards the basal end, which is critical for parasite entry into erythrocyte (Yahata et al, 2021). Glideosome complex, which comprises of several proteins, anchors the actomyosin motor to the inner membrane complex and positions it strategically in the merozoite for its function (Frenal et al, 2017; Frenal et al, 2010).

Protein kinases play a critical role during various cellular processes and signalling events regulated by them are important for most cellular functions. ∼ 85 putative protein kinases and ∼ 30 putative phosphatases are coded in *Plasmodium* genome (Billker et al, 2009; Guttery et al, 2014; Ward et al, 2004). *Plasmodium falciparum* genome wide knockout studies indicated that several of these kinases are refractory to gene disruption suggesting that they are indispensable for parasite survival (Solyakov et al, 2011; Zhang et al, 2018). Several of these kinases are regulated by second messengers like calcium, PIPs, cAMP, cGMP *Plasmodium* kinases like CDPKs, PKA, PKG etc and are crucial for diverse parasitic functions (Alam et al, 2015; Baker et al, 2017a; Bansal et al, 2016; Kumar et al, 2017; Maurya et al, 2022; Perrin et al, 2020). For instance, regulation of cAMP dependent protein kinase PKA by cAMP is critical for host erythrocyte invasion (Perrin et al, 2020; Wilde et al, 2019). PfPKG is involved in egress and invasion of merozoites (Alam et al, 2015; Collins et al, 2013b) and motility of ookinetes (Brochet et al, 2014). cGMP and PKG signalling triggers calcium release from parasite stores which is critical for egress as well as invasion facilitated by the release of microneme proteins (Alam et al, 2015; Collins et al, 2013b; Nofal et al, 2021; Singh & Chitnis, 2012). Calcium plays a critical role in egress and invasion by virtue of its effector kinases like PfCDPK1 (Bansal et al, 2016; Kumar et al, 2017), PfCDPK5 (Dvorin et al, 2010) and phosphatase PfCalcineurin (Paul et al, 2015).

Protein kinase PfPK2, was previously reported to be activated by calcium/calmodulin *in vitro* (Kato et al, 2008) and previous studies also demonstrated that it is indispensable for parasite asexual development (Solyakov et al, 2011; Zhang et al, 2018). However, the precise function of this protein kinase in parasite life cycle has remained unknown. Present studies demonstrate that PfPK2 regulates the invasion of host erythrocyte by *P. falciparum*. It regulates the secretion of AMA1, which is critical for Tight Junction formation. Phosphoproteomics studies revealed that PfPK2 may target proteins that are implicated in invasion, signalling and glideosome assembly. Furthermore, PfPK2 facilitates the formation of cGMP and the release of calcium from intracellular stores, which are critical for host invasion. These and other observations suggested that it may regulate the process of invasion by regulating parasite signalling networks.

## Results

### Recombinant PfPK2 is not activated by calcium/calmodulin *in vitro*

The analysis of PfPK2 (PlasmoDB ID: PF3D7_1238900) suggested that in addition to a kinase/catalytic domain (a.a. 111 to a.a. 364) it has a long N and a C-terminal extensions (Fig. 1A). A previous study suggested that PfPK2 shares reasonable sequence homology with CaM Kinase family members like CamKI (∼33%) (Kato et al, 2008; Zhao et al, 1992) but the sequence similarity is mainly between kinase domains of human CamKIδ and PfPK2 (∼80%) (Kato et al, 2008). PfPK2 has a long N-terminal extension compared to CamKI and a CaM binding domain was predicted within a postulated regulatory domain present downstream of the kinase domain, which is the case with CamKI (Kato et al, 2008). However, a close examination of the sequence in this region suggested a very weak similarity between PfPK2 and CamKIδ in this region (Fig. 1A, B). In addition, PfPK2 also has a long C-terminal extension, which is phosphorylated at several sites in the parasite as indicated by previous phosphoproteomics studies (www.plasmodb.org). Previously, calcium/calmodulin (CaM) was reported to activate PfPK2 *in vitro* (Kato et al, 2008).*(Kato et al, 2008)*

**Figure 1:**
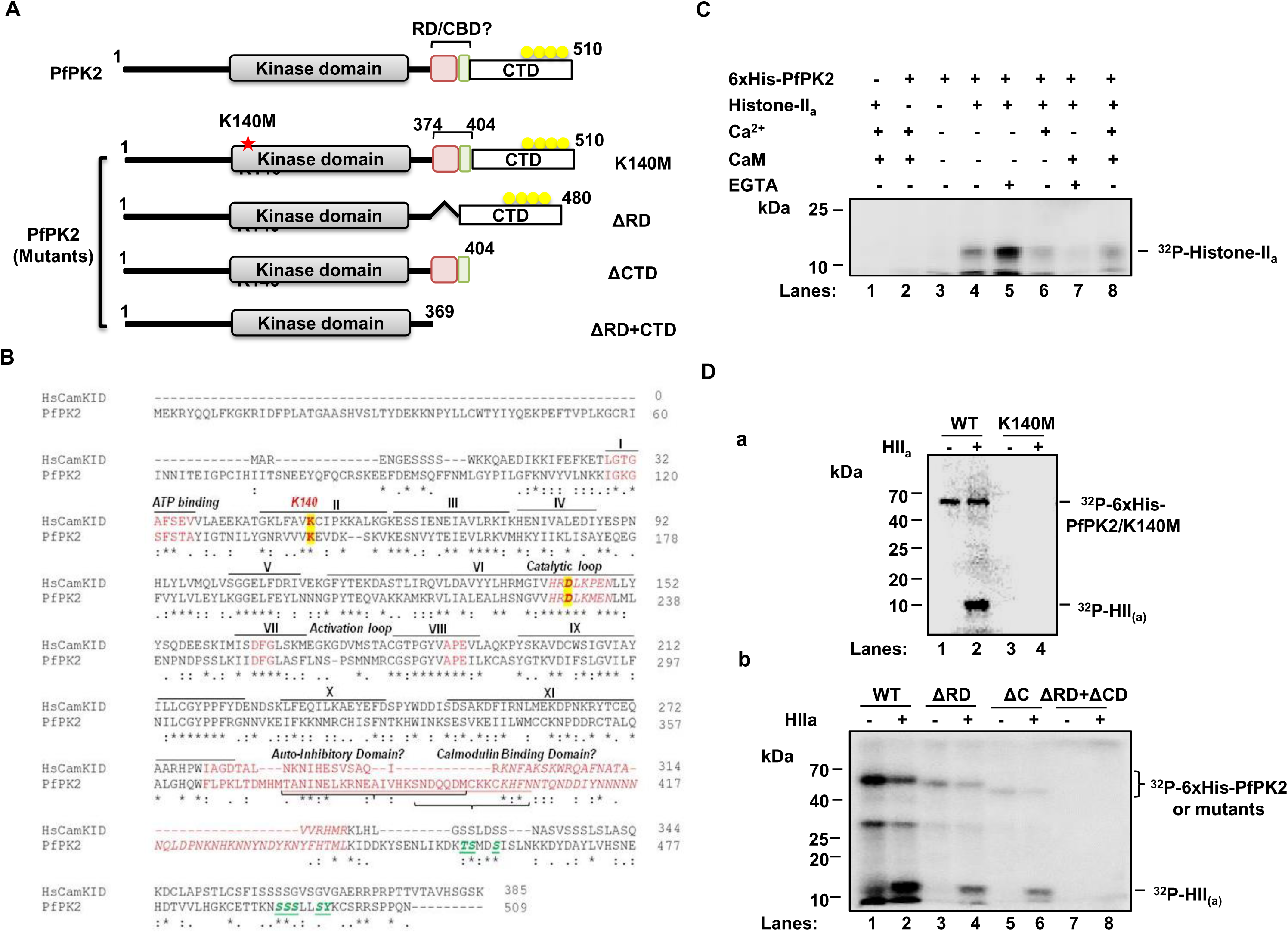
PfPK2 is active *in vitro* independent of calcium/calmodulin. **A.** Schematic indicating domain organization of PfPK2, which includes a putative regulatory domain (RD), a C-terminal domain (CD), which is phosphorylated at several sites in the parasite and a putative calmodulin-binding domain (CBD) predicted by Kato *et al*. (2008). An invariant lysine (K140), which is critical for interaction with ATP in most kinases, is also indicated (red *). Mutants of PfPK2 with K140M mutation or deletion of the RD, CD or RD+CD, which were expressed as recombinant proteins for biochemical analysis, are also indicated. **B.** CLUSTALW alignment of HsCamKID with PfPK2 indicating key motifs, subdomains and critical residues involved in kinase function are highlighted in yellow. The phosphorylation sites identified in the C-terminal domain in the reported phosphoproteomic studies (www.plasmodb.org) are indicated in green. **C.** Purified 6xHis-PfPK2 was used in an *in vitro* kinase assay containing Histone-II_a_ as substrate in a buffer containing [γ^32^P]-ATP in the presence of calcium chloride, calmodulin and/or EGTA as shown in the figure. After SDS-PAGE, phosphorimaging was performed to detect the phosphorylation of the indicated proteins. **D.** Kinase assay was performed using equal amount of purified 6xHis-PfPK2 or its K140M (a), ΔRD, ΔCD, ΔRD+ ΔCD mutants (b) as described in panel A.

Recombinant PfPK2 was expressed as a 6xHis-tagged protein in *E. coli* for biochemical characterization. A kinase assay was performed in which Histone II_a_ was used as a phosphoacceptor substrate. Recombinant PfPK2 phosphorylated histone II_a_ (Fig. 1C). However, there was no increase in its activity upon addition of calcium/calmodulin. In contrast, there was a slight decrease in histone phosphorylation when calcium/calmodulin was added to the reaction mix (lane 6/7 vs lane 5). These data suggested that PfPK2 may not be activated by calcium/calmodulin, at least *in vitro*.

A lysine at 140 position (K140) (Fig. 1B), which is complementary to an invariant lysine present in subdomain II of most PKs and is critical for their activity (Iyer et al, 2005), was mutated to M (K140M). K140M mutant did not exhibit any autophosphorylation or Histone phosphorylation (Fig. 1Da), which suggested that this residue is critical for PfPK2 activity. Next, we analyzed the role of the C-terminal domain (CD) and putative regulatory domain (RD) in PfPK2 regulation. For this purpose, deletion mutants lacking these domains were generated and recombinant proteins were used for activity assays. The deletion of RD or CD caused a decrease in the phosphorylation of Histone II_a_ by PfPK2 (Fig. 1Db). A marked decrease in autophosphorylation was also observed in the case of ΔRD, which was even more significant in the case of the ΔCD mutant. The deletion of ΔRD and ΔCD resulted in almost complete loss of PfPK2 activity. Collectively, these data suggested that at least *in vitro* these domains are critical for PfPK2 activity. It is possible that the phosphorylation of CD, which is also phosphorylated at several sites in the parasite (www.plasmodb.org), may have a role in its parasitic functions like interaction with other proteins.

### Role of PfPK2 in the development of *P. falciparum*

#### Conditional knockdown of PfPK2 in P. falciparum

Previous attempts to knockout PfPK2 from *P. falciparum* have been unsuccessful suggesting that this kinase may be indispensable for blood stage development of the parasite (Solyakov et al, 2011). However, its role in *P. falciparum* remained undefined. We used a rapamycin-inducible dimerizable Cre recombinase (diCre) based strategy for conditional knockdown of PfPK2 and used 1G5DC strain (Collins et al, 2013a), which expresses FKBP and FBP that are independently fused to two halves of the Cre recombinase and dimerize upon addition of rapamycin (RAP) resulting in active Cre enzyme, which can then facilitate the excision of the genetic sequence flanked by two loxP sites. Using Selection Linked Integration (SLI) (Birnbaum et al, 2017) loxP sites were introduced in the PfPK2 locus (Fig. 2A) and a recodonized copy of PfPK2 was introduced downstream of GFP. Parasites obtained after drug (WR99210 and DSM1) selection were subjected to limiting dilution cloning. Correct integration of the targeting plasmid at the desired locus was confirmed by PCR as amplicons of expected size were obtained parasites from a clone that lacked wild type PfPK2 (Fig. 2B). The addition of rapamycin (RAP) to PfPK2-loxP parasites for 4h or 24h at ring stages resulted in efficient excision of the floxed sequence at the schizont stage of the same cycle (Fig. 2C). Successful expression of GFP-PfPK2 was indicated by Western blotting (Fig. 2D) as well as by IFA (Supp. Fig. S1A), which wasundetectable after rapamycin treatment suggesting efficient depletion of this kinase from the parasite (Fig. 2D and Supp. Fig. S1B). IFA indicated that PfPK2 localized to punctate vesicular structures often present apically near the micronemes but did not exhibit any significant co-localization with microneme (AMA1) or rhoptry (RhopH3) proteins (Supp. Fig. 1A).

**Figure 2:**
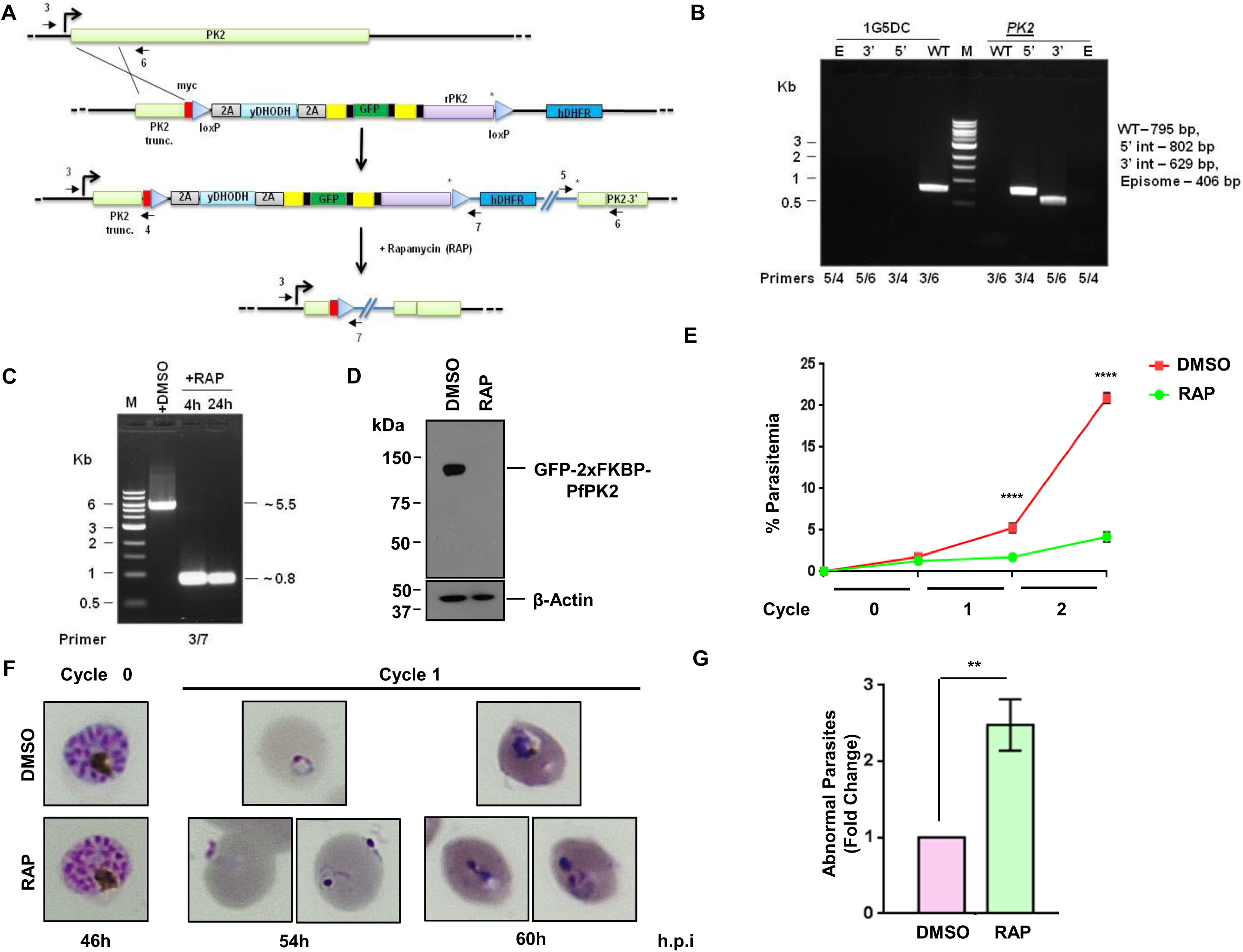
PfPK2 is critical for asexual phase life cycle of *P. falciparum*. **A.** Schematic representation of the strategy used to modify the *PfPK2* locus through single-crossover homologous recombination and selection-linked integration (SLI). A 330bp region homologous to the 5’-end of *PfPK2* gene was cloned in pSLI-N-sandwich-loxP vector. In addition, recodonized PfPK2 (rPK2) was cloned in frame with GFP (green). Yellow-FKBP, black-linker, triangles -loxP, *-stop codon, arrows-location of annealing site of PCR primers used for genotyping and assessing excision, which was achieved after rapamycin addition. **B.** Genotyping of a clone obtained after limited dilution (PfPK2-loxP). PCR amplification was carried out for both the unmodified and modified loci, using primers targeting the indicated sites in Panel A. PCR products of expected size, which is indicated in the figure, were amplified for assessing 5’ and 3’-integration, presence of wild type (WT) locus or episome (E) was absent in the transgenic PfPK2-loxP parasite. The amplicons were sequenced to further confirm the integration at the desired locus. **C.** PfPK2-loxP parasites were treated with DMSO or 200 nM RAP for 4 or 24h. Subsequently, genomic DNA was isolated ∼44 hpi and PCR was performed using indicated primer sets (panel A). A ∼0.8kb product was obtained upon RAP treatment indicating successful excision. **D.** Western blot examination of PfPK2-loxP lysates treated with DMSO and RAP and using anti-GFP antibody revealed that the RAP-treatment resulted in effective depletion of GFP-PfPK2. As a loading control, a β-actin antibody was also used to probe the blots. **E.** PfPK2-loxP parasites were synchronized and ring stage parasites were used for setting up growth rate assay in the presence of DMSO or RAP. After 6h, RAP was washed and parasite growth was assessed after each cycle by performing flow cytometry (SEM ± SE, n=3, ANOVA, **** P<0.0001; ns -not significant). **F.** PfPK2-loxP parasites were treated with DMSO or RAP as described in panel A and Giemsa-stained blood smears at indicated time post invasion (h.p.i) were examined. PfPK2- depleted parasites showed significant parasites attached to the erythrocytes were suggestive of failed invasion. A few parasites that entered erythrocytes exhibited abnormal morphology and/or were pyknotic. **G.** The number of abnormal/pyknotic parasites -indicated in panel G-were counted after RAP addition (72 h.p.i), which revealed a significant increase in number of abnormal/pyknotic RAP-treated parasites when compared to DMSO-treated controls (SEM ± SE, n=3, ANOVA, ** P<0.01).

#### PfPK2 is important for erythrocyte invasion and asexual development of P. falciparum

To investigate the role of PfPK2 in parasite development, synchronized ring stage cultures of 1G5DC (control) or PfPK2-loxP lines were treated with RAP to promote the excision of PfPK2. PfPK2-loxP parasites did not exhibit much change in parasitemia in the cycle of RAP-treatment (cycle 0). However, the parasite growth was significantly reduced at the onset of the next cycle and very few parasites were observed subsequently (Fig. 2E). There was almost no apparent change in number of merozoites formed upon RAP-treatment during cycle 0 (Supp. Fig. S2B) suggesting parasite replication was unaffected.

A close examination of Giemsa-stained blood smears of parasite cultures at the beginning of cycle 1 revealed a very small number of ring-infected erythrocytes upon PfPK2 depletion, which was indicative of possible defects in egress and/or invasion (discussed below in detail). A very small number of parasites that managed to invade exhibited stunted abnormal morphology and/or were pyknotic and failed to mature to trophozoites (Fig. 2F and 2G).

The possibility of defects in egress and/or invasion upon PfPK2 depletion was investigated in detail. For this purpose, assays were performed using DMSO or RAP-treated mature schizonts (cycle 0, ∼40-44 hpi), which were incubated with fresh erythrocytes. The schizont number consistently reduced with almost no residual schizonts left after a few hours, which suggested that the egress was not impaired in PfPK2-depleted parasites (Fig. 3A). In contrast, there was a marked decrease in the number of fresh rings formed in RAP-treated parasites (Fig. 3A and 3B), which was indicative of impaired invasion. A parasite line was generated in PfPK2-loxP background in which a WT copy of HA-tagged PfPK2 was expressed episomally to complement the PfPK2 function (cHA-PfPK2-loxP) as RAP does not influence the expression of HA-PfPK2 (Supp. Fig. S2A). Importantly, there was no significant difference in invasion as well as post-invasion defects reported above in the case of cPfPK2-loxP parasites (Supp. Fig. S2C and S2D) confirming the involvement of PfPK2 in invasion and in these processes.

**Figure 3:**
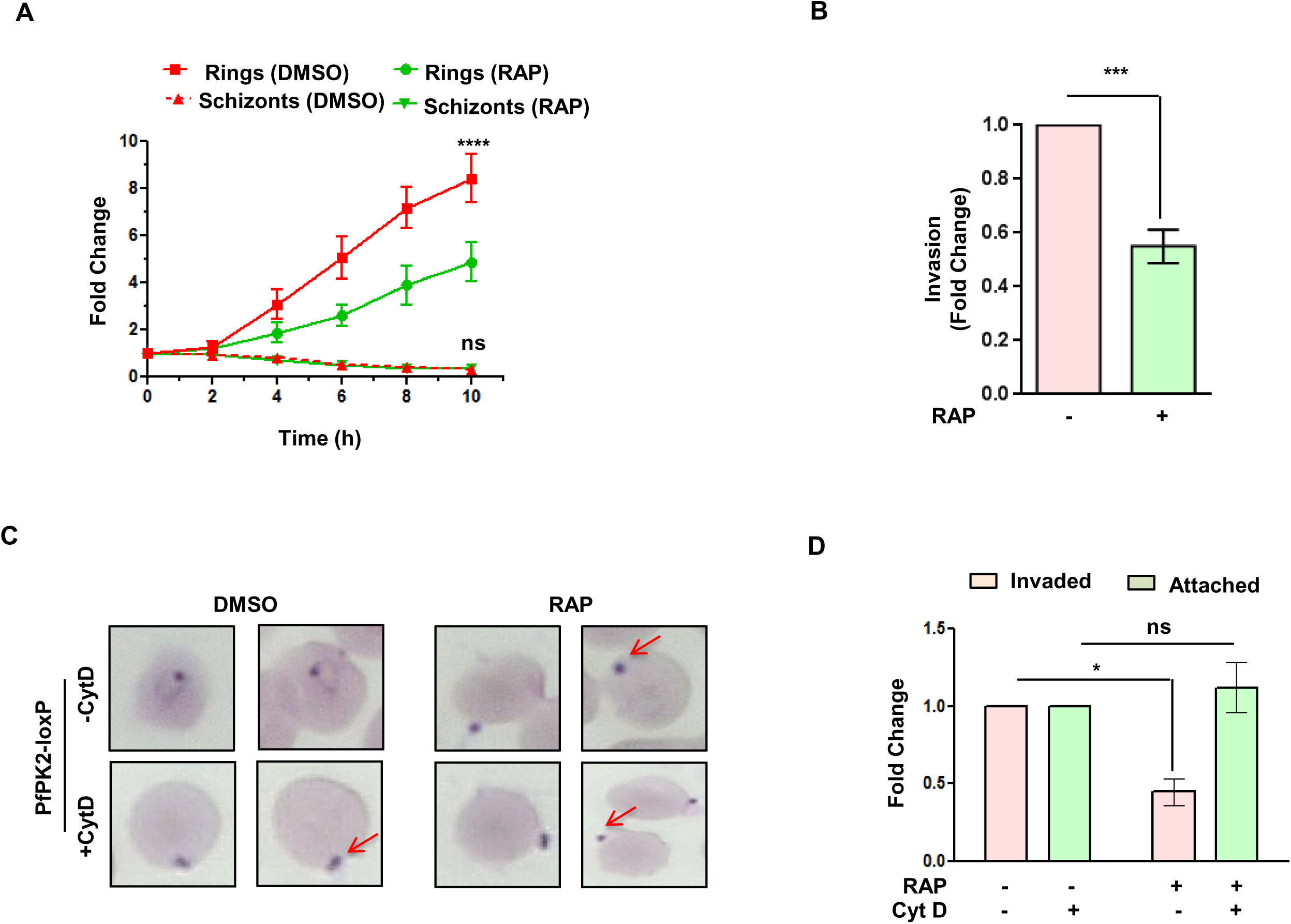
PfPK2 regulates the process of host erythrocyte invasion. **A.** PfPK2-loxP schizonts treated with DMSO or RAP were incubated with fresh erythrocytes. After indicated time, samples were collected and the number of schizonts and rings were counted from Giemsa-stained thin blood smears and fold change in schizont or ring- infected erythrocytes was determined. RAP-treated parasites showed significant decrease in the number of rings whereas schizont numbers were largely unaltered and almost no schizonts were observed at the end of the assay (SEM ± SE, n=6, ANOVA, **** P<0.0001). **B.** PfPK2-loxP schizonts were treated with DMSO or RAP and were allowed to invade fresh erythrocytes as indicated in panel A. Fold change in ring-stage parasitemia, which reflected invasion, was determined after ∼12 h.p.i from experiments described in panel A (SEM ± SE, n=3, ANOVA, *** P<0.001). **C and D.** DMSO or RAP treated PfPK2-loxP parasites were used for erythrocyte attachment assays which were performed in the presence or absence of cytochalasin D. The number of merozoites attached to erythrocytes were counted from Giemsa-stained thin blood smears (C) and fold change in % attachment (D) with respect to DMSO treated parasites was determined (SEM ± SE, n=3, ANOVA, * P<0.05; ns -not significant).

Further investigations were performed to determine which stage of host erythrocyte invasion, which is a complex multi-step process (Duraisingh et al, 2008; Weiss et al, 2015), is regulated by PfPK2. Upon release after the rupture of erythrocytes, merozoites first attach to the surface of fresh erythrocytes and several ligand-receptor interactions are involved in this process. The role of PfPK2 in attachment was probed by using cytochalasin D (Cyt D), which prevents invasion or entry but does not affect attachment (Miller et al, 1979). There was no significant difference in the number of DMSO or RAP-treated PfPK2-loxP parasites attached to erythrocytes (Fig. 3C and 3D), which suggested that PfK2 may regulate an event post-attachment.

Live cell imaging was performed for a more detailed examination of defects in the invasion process as a result of PfPK2 depletion. Typically, merozoite invasion starts with the attachment of the merozoite to the host erythrocyte resulting in the deformation of the erythrocyte membrane (Weiss et al, 2015). Subsequently, the merozoite reorients its apical end on to the erythrocyte surface followed by its penetration into the erythrocyte. Once internalized, the infected erythrocyte undergoes echinocytosis for a short period followed by its recovery. Subsequently, erythrocyte membrane seals and the parasite attains ring like morphology (Fig. 4A,a Video S1) (Geoghegan et al, 2021; Weiss et al, 2015). Video microscopy revealed significantly lesser RAP-treated merozoites invading the erythrocytes in comparison to DMSO-treated counterparts (Fig. 4, Video S1-S4,). Upon RAP-treatment most parasites remained attached to the erythrocyte surface and also exhibited re-orientation with apical end positioned to enter the erythrocyte (Fig.4A,b,Video S2, Panel C). The attachment was prolonged but parasites failed to penetrate the erythrocyte. The attached merozoites were able to deform the host erythrocyte membrane and caused echinocytosis for a significantly longer duration in RAP- treated parasite (Fig. 4A, panel c and d, Video S3-S4, Panel C). These observations suggested that PfPK2 depletion may not influence attachment but it prevents entry possibly at a late step after interaction between the merozoite and the erythrocyte surface, which was supported by the fact that attached parasites caused deformation as well as echinocytosis. Collectively, these observations suggested a role of PfPK2 in later-steps of invasion like formation of Tight Junction and possibly subsequent sealing of the erythrocyte.

**Figure 4:**
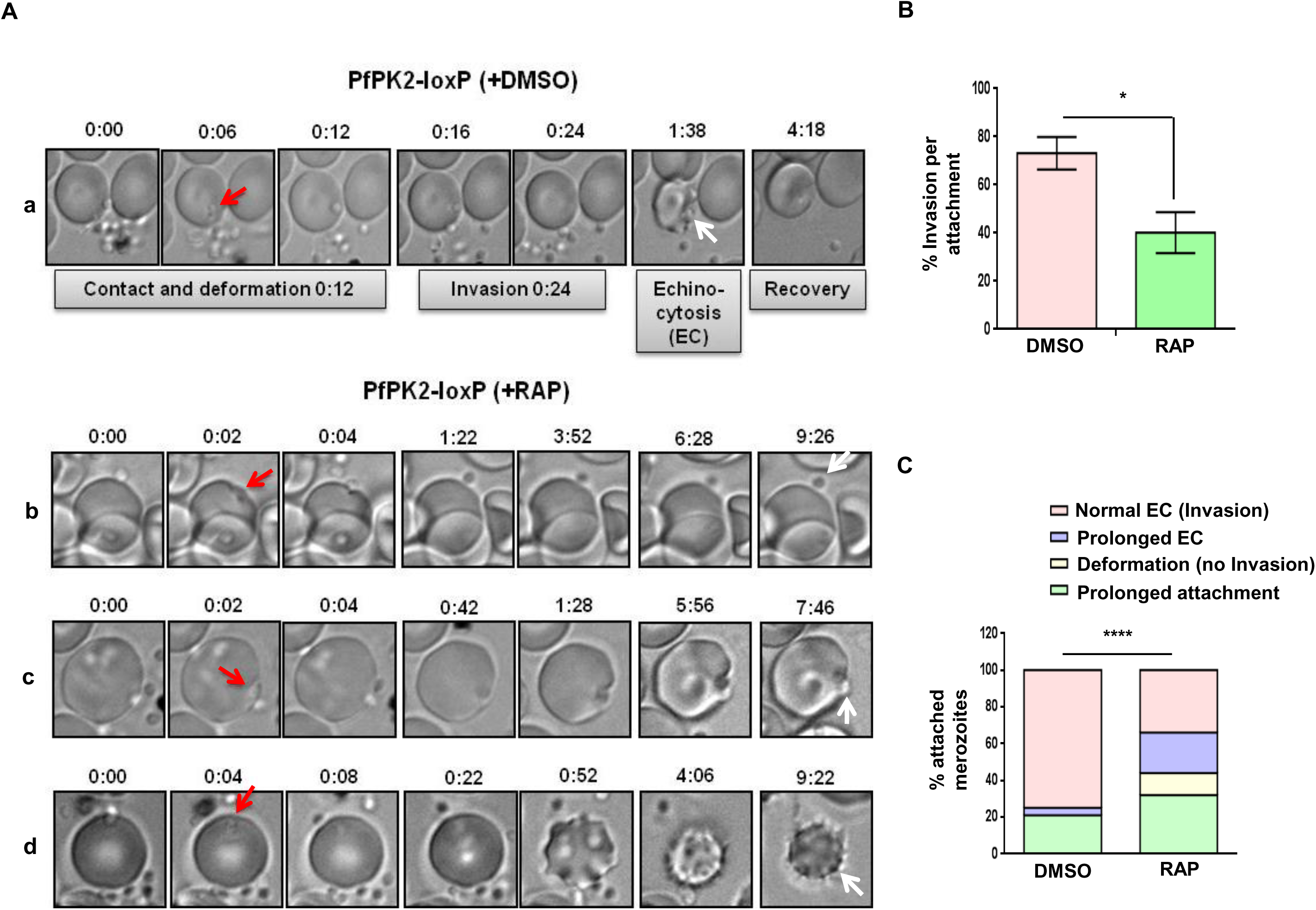
PfPK2 depletion impairs the entry of merozoites in host erythrocytes. **A.** Schizonts isolated from PfPK2-loxP parasites cultured in the presence or absence (DMSO) of RAP for one cycle and post-egress merozoites were used for live microscopy (Video S1-S4). Selected still images from videos are used to illustrate various stages of invasion. DMSO-treated PfPK2-loxP merozoites successfully deform the erythrocyte membrane (red arrow), invade, and trigger echinocytosis (white arrow) followed by recovery of the infected erythrocyte (a, video S1). Prolonged attachment upon RAP-treatment and unsuccessful invasion was observed (b, video S2). Several RAP-treated parasites that exhibited prolonged attachment (red arrow) were also able to deform the erythrocyte membrane (c, video S3) and stimulate echinocytosis (EC) which was for much longer duration (d, white arrow, video S4). **B.** % invasion per attachment was determined and data represent results from four independent pooled experiments described in panel A (mean ± SE, n = 4, *, P < 0.05, paired t test). **C.** Various outcomes during erythrocyte invasion indicated in Panel A were quantitated for merozoites that formed an initial attachment using data from four different experiments for each set of conditions [****, P < 0.0001, chi-square test].

#### PfPK2 regulates the discharge and shedding of AMA1 from the parasite

As mentioned above, discharge of ligands from secretory organelles like micronemes and rhoptries is critical for invasion (Cowman et al, 2017; Duraisingh et al, 2008). Therefore, we assessed the release of rhoptry and microneme proteins in parasite culture supernatants. There was almost no difference in the release of rhoptry protein RhopH3 upon RAP treatment (Fig. 5A, Supp. Fig. S3C) excluding the possibility of involvement of PfPK2 in rhoptry release. Interestingly, the shedding of microneme protein AMA1 - which is involved in Tight Junction (TJ) formation (Giovannini et al, 2011; Srinivasan et al, 2011; Treeck et al, 2009; Yap et al, 2014) – was dramatically reduced upon PfPK2 depletion (Fig. 5A, Supp. Fig. S3A). The release of another microneme protein EBA-175, which interacts with glycophorin A on the surface of the erythrocytes (Sim et al, 1994), was also reduced (Fig. 5A, Supp. Fig. S3B). but its localization to the microneme was unaltered (Supp. Fig. S4). These data suggested that PfPK2 may regulate microneme release. However, there was no difference in the attachment of parasites (Fig. 3C), which is regulated by adhesins like EBA1-75. Therefore, we focussed on a possible role of PfPK2 in the release of AMA1, which is involved in late inavsion events like Tight Junction formation and PfPK2 plays a role at this stage of invasion (Fig. 4). AMA-1 is first secreted to the merozoite surface from micronemes and subsequently shed or released by the action of proteases like SUB2 and rhomboid proteases (Dutta et al, 2003). AMA1 was found in micronemes as indicated by a puncta at the apical end in both DMSO or RAP- treated parasites but a significant population was found on merozoite surface. In contrast, there was a decrease in parasites with AMA-1 on their surface upon PfPK2 depletion and it was confined mainly to the micronemes (Fig. 5B). Therefore, it appears PfPK2 may regulate the secretion of AMA-1 on to the merozoite surface and in its shedding from merozoite surface.

**Figure 5:**
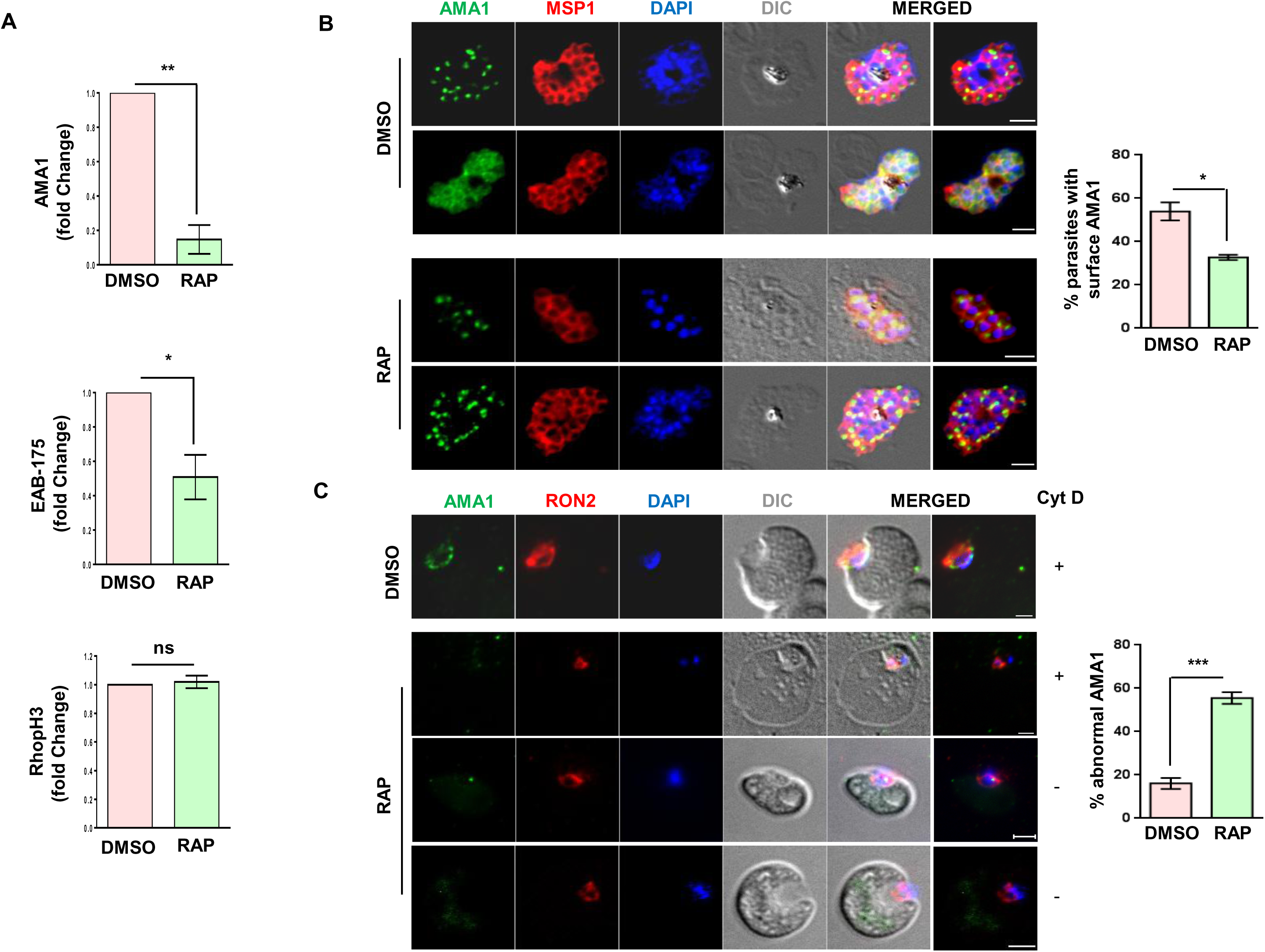
PfPK2 regulates microneme release. **A.** DMSO or RAP treated PfPK2-loxP schizonts were cultured for one cycle and the release of AMA1, EBA175 and RhopH3 from parasites was determined by performing Western blotting on culture supernatant using specific antibodies (Supp. Fig. S3). The secretion of these proteins in the supernatant was quantitated by densitometry of the Western blot, and the fold change in secretion for RAP-treated parasites with respect to DMSO-treated parasites was determined (AMA1, mean ± SEM, n = 4, **, P < 0.01; EBA175, mean ± SEM, n = 4, *, P < 0.05; RhopH3, mean ± SEM, n = 4, ns, P > 0.05, ns-not significant; paired t test). Western blots were also performed on total parasite protein lysates, which were used for normalization of secreted proteins (Supp. Fig. S3). **B.** PfPK2-loxP parasites were treated with DMSO or RAP. Subsequently, schizonts were treated with E64 to prevent egress and IFA was performed to detect AMA1 and MSP1. AMA1 localizes to microneme in schizonts stage (DMSO, upper panel) but during egress it comes to the merozoite surface in a significant number of parasites (DMSO, lower panel). In RAP-treated parasite AMA1 secretion to parasite surface was reduced as it was retained in the micronemes in most parasites, which was confirmed by counting parasites with AMA1 staining on the surface (mean ± SEM, n=3, * p<0.05, t test). **C.** The presence of AMA1 at the Junction in DMSO or RAP-treated PfPK2-loxP parasites was determined by performing IFA for AMA1 and RON2. In the case of DMSO-treated parasites, AMA1 localized proximal to RON2 whereas it was either not detected or it was significantly reduced in RAP-treated parasites. *Right panel*, parasites exhibiting abnormal or no AMA1 staining were quantitated [SEM ± SE, n=3, ANOVA, *** P<0.001)].

AMA1 plays a role in the formation of Tight-Junction (TJ) is *via* its association with rhoptry neck proteins (RONs) that are released by the rhoptries “Just-in-Time” on to erythrocyte surface (Riglar et al, 2011). IFAs were performed to localize AMA1 along with RON2 in “invading” merozoites. As reported previously (Baum et al, 2017; Riglar et al, 2011; Riglar et al, 2016), AMA1 either forms a ring around the surface and/or was concentrated at the points of contact between the merozoite and the erythrocyte membrane along with RON2 (Fig. 5C). RAP-treatment caused a dramatic decrease in AMA1 staining, which was either not observed or at best was seen as a small puncta that did not co-localize with RON2 (Fig. 5C). These data were consistent with the fact that AMA1 secretion was impaired upon PfPK2 depletion (Fig. 5A). Given that AMA1 is critical for TJ formation (Yap et al, 2014), it is reasonable to state that PfPK2 may contribute to this process.

#### Identification of PfPK2 targets by quantitative phosphoproteomics

In order to elucidate the mechanism via which PfPK2 may regulate parasite invasion, it was important to identify the targets of PfPK2 and unravel PfPK2-dependent signalling pathways. Therefore, quantitative phosphoproteomics was used to compare the phosphoproteome of PfPK2-loxP depleted parasites in the presence or absence of RAP (Fig. 6A). PfPK2 expression was maximal in late schizonts/merozoites during asexual blood stage development, which correlates well with defects in invasion (Fig. 4). Therefore, it was pertinent to use late schizonts for phosphoproteomics. The lysates of RAP or DMSO-treated PfPK2-loxP parasites were harvested at the late schizont stage, and phosphoproteomic analysis was performed on the prepared lysates. Phosphoproteomics identified 2894 phosphopeptides in the PfPK2-loxP parasites and comparison of the phoshoproteome of DMSO and RAP-treated parasites indicated that several sites on these peptides exhibited differential phosphorylation upon PfPK2 knockdown achieved by RAP treatment (Fig. 6B). A total of 104 phosphosites on 82 proteins exhibited significant reduction in phosphorylation upon PfPK2 depletion (Fig. 6A, Supp Data Set S1). It was interesting to note that several proteins with well-defined or proposed functions in invasion exhibited significant changes in phosphorylation, including proteins involved in signalling pathways and actomyosin motor function (Fig. 6B, Table 1). Strikingly, several key proteins were also hyperphosphorylated in PfPK2-depleted parasites, which may be due to direct or indirect effects exerted by PfPK2 on a protein phosphatase or by negatively regulating other protein kinases. For instance, protein phosphatase PPM2 was found to be hyperphosphorylated upon PfPK2 depletion, which may potentially contribute to the phosphorylation status of some of the aberrantly phosphorylated proteins.

**Figure 6:**
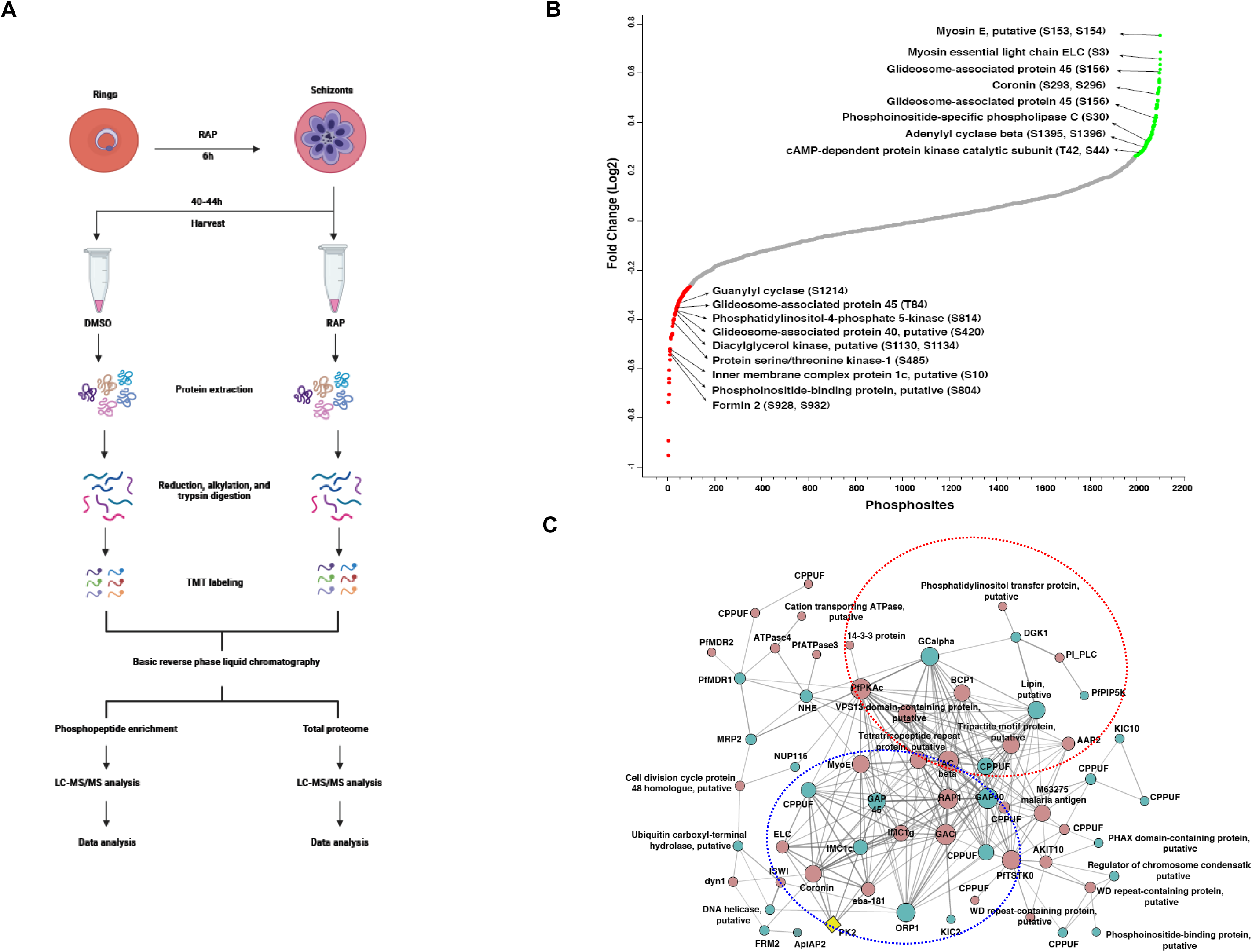
Identification of PfPK2 targets in the parasite by comparative phosphoproteomics. **A.** PfPK2-loxP parasites were synchronized to obtain rings, which were treated with DMSO (control) or RAP and parasites were harvested at schizont stage to perform comparative phosphoproteomic and proteomic analyses as indicated in the schematic (See Methods for details). **B.** The phosphorylation fold-change ratios of identified phosphopeptides were normalized to total protein abundance fold-change. The fold-change ratios for all phosphopeptides from various replicates are provided in Supplementary Data Set S1.1. The S-curve for the normalized data is provided for some of the significantly altered phosphorylation sites belonging to key proteins (Supplementary Data Set 1.2, 1.3) are indicated. **C.** Protein-protein interactions were predicted between differentially phosphorylated proteins using the STRING resource. The analysis exhibited high confidence interactions between the candidate proteins (Supplementary Data Set 1.5). Two major protein-protein interaction clusters involving proteins implicated in signalling and glideosome/invasion (Table 1) are encircled in red and blue, respectively. Red and green nodes illustrate hyperphosphorylated and hypophosphorylated proteins, respectively. The term “CPPUF” refers to a Conserved *Plasmodium* Protein with an unknown function.

**Table 1.** Putative parasitic targets of PfPK2 that is implicated in invasion and signalling. Significantly altered hyper- and hypo-phosphorylated sites are indicated in pink and green boxes, respectively.

| Phosphorylated proteins | PlasmoDB ID | Phosphosites |
| --- | --- | --- |
| **Signalling** |  |  |
| Adenylyl cyclase $\beta$ | PF3D7_0802600 | S1395, S1396 |
| Protein phosphatase PPM2 | PF3D7_1138500 | S139, T142 |
| cAMP-dependent protein kinase catalytic subunit | PF3D7_0934800 | T42, S44 |
| PI-Phospholipase C | PF3D7_1013500 | S30 |
| Phosphatidylinositol-4-phosphate 5-kinase | PF3D7_0110600 | S814 |
| Diacylglycerol kinase, putative | PF3D7_0930500 | S1130, S1134 |
| Guanylyl cyclase $\alpha$ | PF3D7_1138400 | S1214 |
| **Invasion/Egress** |  |  |
| Erythrocyte binding antigen-181 | PF3D7_0102500 | S1299 |
| Rhoptry-associated protein 1 | PF3D7_1410400 | S140, S169, S294 |
| Merozoite surface protein 2 | PF3D7_0206800 | S214 |
| Rhoptry-associated protein 1 | PF3D7_1410400 | S127 |
| **Glideosome/IMC** |  |  |
| Coronin | PF3D7_1251200 | S293, S296 |
| Inner membrane complex protein 1c, putative | PF3D7_1003600 | S266, S267 |
| Inner membrane complex protein 1g, putative | PF3D7_0525800 | S277 |
| Myosin E, putative | PF3D7_0613900 | S153, S154 |
| Myosin essential light chain (ELC) | PF3D7_1017500 | S3 |
| Glideosome-associated protein (GAP) 45 | PF3D7_1222700 | S156 |
| Glideosome-associated connector (GAC) | PF3D7_1361800 | T564, S566 |
| Glideosome-associated protein 40, putative | PF3D7_0515700 | S420 |
| Inner membrane complex protein 1c, putative | PF3D7_1003600 | S10 |
| Glideosome-associated protein 45 | PF3D7_1222700 | T84 |

The above-mentioned family of proteins were of special interest as they are likely to be relevant for the function of PfPK2 in host erythrocyte invasion. Glideosome associated proteins PfGAP45 and PfGAP40 exhibited altered phosphorylation upon PfPK2 depletion. These proteins are important for anchoring the motor and thereby regulate actomyosin motor activity (Frenal et al, 2017; Keeley & Soldati, 2004), which is essential for erythrocyte invasion (Perrin et al, 2018). Previously, phosphorylation of PfGAP45 was reported by several studies which is considered to be important for invasion (Ridzuan et al, 2012; Thomas et al, 2012).

The catalytic subunit of cAMP-dependent protein kinase PfPKA exhibited hyperphosphorylation upon PfPK2 depletion. Strikingly, PfPKAc hyperphosphorylation was observed at T42 and S44 (Fig. S6B, Table 1, Supp Dataset S1) which coincides with the ATP binding region that comprises of a GxGxxG motif. The phosphorylation of these sites may have a bearing on interaction of ATP with the kinase and inhibit the activity of PfPKA (Hemmer et al, 1997; Steinberg, 2018). In addition, adenylyl cyclase (ACβ) was also found to be hyper phosphorylated. These observations hinted at possible involvement of the cAMP signalling module downstream of PfPK2 and altered phosphorylation of these proteins can have a bearing on cAMP signalling and PfPKA activity. Interestingly, PfPKA has been shown to play a role in erythrocyte invasion by the parasite by several groups (Leykauf et al, 2010; Perrin et al, 2020; Wilde et al, 2019).

Strikingly, PfPK2 depletion prevented phosphorylation of gunaylyl cyclase PfGCα (Fig. 6B, Table 1), which is involved in generating second messenger cGMP. cGMP signalling regulates host erythrocyte invasion as well as egress (Alam et al, 2015; Collins et al, 2013b; Moon et al, 2009; Nofal et al, 2021). Several proteins implicated in phosphoinositide (PIP) metabolism and or downstream effectors of PIP signalling were aberrantly phosphorylated upon PfPK2 depletion: A PI4P-5-Kinase, which is involved in generation of PI(4,5)P2, was hypophosphorylated at S814 (Fig. 6B, Table1, Supp Data Set S1).

PI4,5P2 can regulate signalling via downstream effector proteins and/or serve as substrate for PI-PLC. Interestingly, PfPLC phosphorylation at S30 increased in PfPK2-depleted parasites (Fig. 6B,Table 1, Supp Data Set 1). PLC hydrolyses PI(4,5)P2 to yield I(1,4,5P3) (IP3) and diacylglycerol (DAG), which are potent second messengers. IP3 regulates calcium release from the ER in the parasite (Vaid et al, 2008a) like other organisms but IP3 receptor has not been identified, although a transmembrane protein has been implicated in this process (Balestra et al, 2021). PLC-mediated calcium release has been implicated in the process of invasion (Vaid et al, 2008a). DAG is substrate for DAG-kinase, which converts it to phosphatidic acid (PA). Recently, DAGK inhibition was shown to block gliding motility which is important for invasion (Yahata et al, 2021). Therefore, it was interesting to note that the phosphorylation at S1130 and S1134 was reduced when PfPK2 was depleted (Fig. 6B, Table 1, Supp Data Set 1). Given that these pathways are involved in invasion, it is reasonable to suggest that PfPK2 may regulate invasion by possibly regulating cGMP and/or phosphoinositide signalling in the parasite.

Protein-protein interaction (PPI) networks among differentially phosphorylated proteins (Fig. 6C) were predicted using the STRING database, which uses information from reported experimental results, co-expression profiles etc (Szklarczyk et al, 2015). Consistent with observations made above, two major PPI- modules that related to PfPK2 function in invasion emerged which comprised of: a. proteins related to invasion like surface, rhoptry and microneme proteins (e.g. RAP1, EBA181) or gideosome/actomyosin motor related proteins (GAP45, GAC, GAP40, MyoE) ; b. signalling proteins like GCα, PKAc, ACβ, PLC (Fig. 7C). These putative PPI-networks provided first indication of the molecular processes possibly regulated by PfPK2 during the process of invasion.

**Figure 7:**
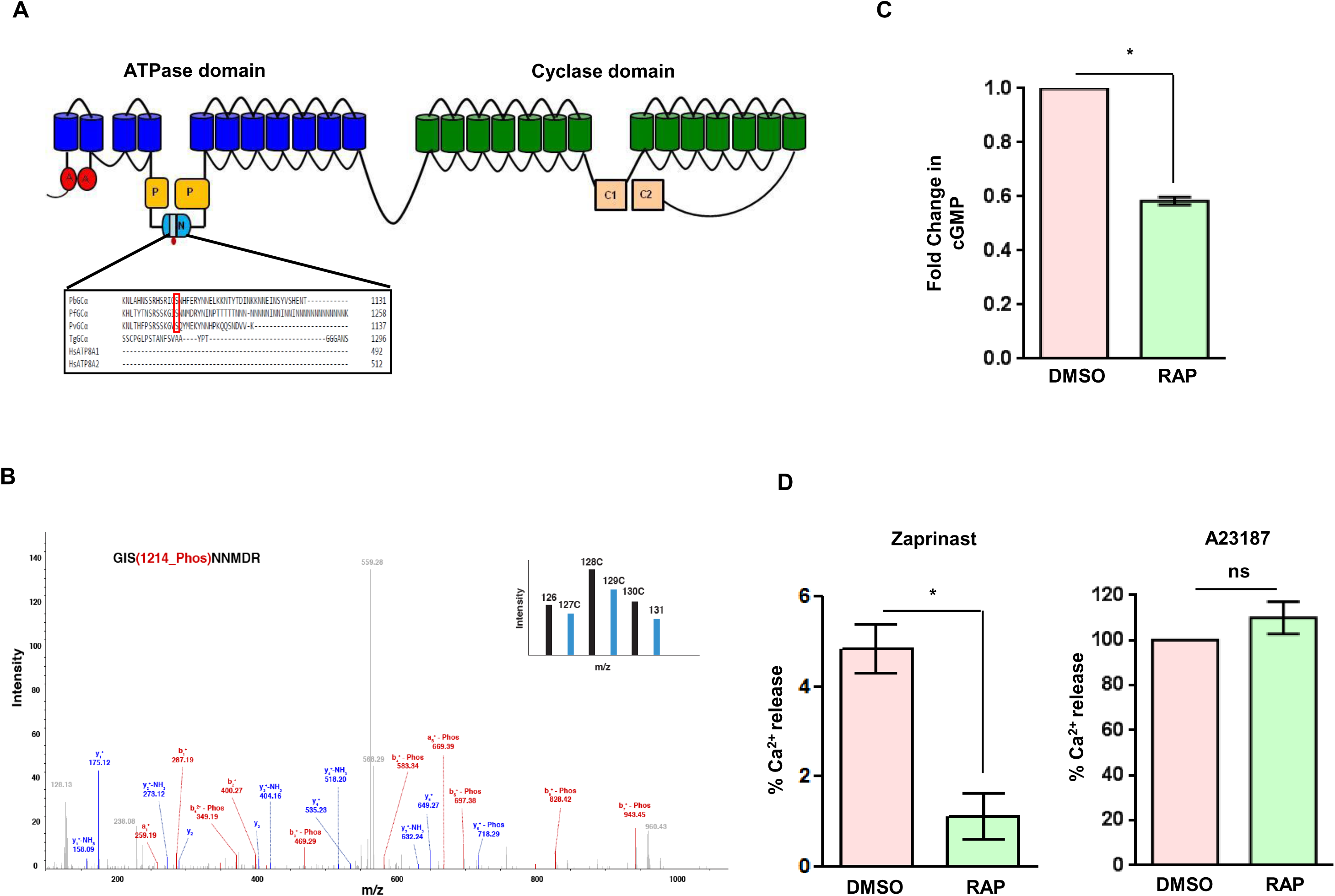
PfPK2 regulates calcium release and cGMP levels in the parasite. **A.** Schematic illustrating the domain architecture of PfGCα, which has an ATPase domain fused to the guanylyl cyclase, various subdomains are indicated (Nofal et al, 2021). The Nucleotide binding domain (N), which splits the phosphorylation (P) domain, has a long insert and PfPK2-regulated phosphorylation site S1214 resides in an insert. Sequence alignment of GCα from *P. falciparum, P. berghei and Toxoplasma gondii* and human ATPase ATP8A1/2 indicates that this site is conserved only in *Plasmodium* spp. **B.** The phosphorylation of S1214 of PfGCα was reduced upon PfPK2 depletion. MS/MS spectra is provided for the corresponding phosphopeptide. TMT channel indicated in the inset represents independent biological replicates of PfPK2 DMSO (black) or RAP (cyan) parasites and wild type samples. **C.** DMSO or RAP-treated PfPK2-loxP schizonts were lysed and a ELISA-based assay kit was used to quantitate intracellular cGMP. The data provides fold change in cGMP upon RAP-treatment (mean ± SEM, n = 3, *, P < 0.05, ANOVA). **D.** Fluorimetry was used to assess calcium mobilization, which was determined by using Fluo-4-loaded mature DMSO- and RAP-treated PfPK2-loxP schizonts that were treated with zaprinast (left panel) or A23187 (right panel). The signals were normalised with respect to DMSO (0%) and A23187 (100%). The data presented are mean of four independent biological replicates that were done in duplicate (mean ± SEM, n = 4, *, P < 0.05, ns, P > 0.05, ns- not significant, paired t test).

#### PfPK2 regulates cGMP levels and calcium release in the parasite

As described above, phosphoproteomic studies indicated that PfPK2 depletion altered the phosphorylation of several signalling proteins that includes guanylyl cyclase GCα coded by gene PF3D7_0381400 (Fig. 6B, Table1 and Fig. 7A,B) which synthesizes cGMP in the parasite (Nofal et al, 2021). In the case of *Plasmodium spp*., GCs are assembled in a peculiar configuration in which the cyclase domain comprising of the two catalytic domain is fused to a P4-ATPase domain (Baker et al, 2017a). The nucleotide binding domain (N-domain), which is important for ATP binding, has large inserts in the case of PfGCα (Nofal et al, 2021). Interestingly, phosphorylation site S1214 which is regulated by PfPK2 (Fig. 6B, 7B) resides in this insert This phosphositeis conserved only in GCα from *Plasmodium* spp (Fig. 7A) and is absent from human ATPase8A1/2.

To explore if PfPK2 depletion has an impact on cGMP synthesis, levels of this cyclic nucleotide were measured in the parasite. A phosphodiesterase (PDE) inhibitor zaparinast, which prevents cGMP degradation, was used to maintain cGMP levels for detection. Strikingly, PfPK2-depletion caused a marked decrease in cGMP formation upon PfPK2 depletion indicating its role in the synthesis of this second messenger (Fig. 7C).

Given that calcium release in *P. falciparum* is regulated by cGMP (Nofal et al, 2021), it was pertinent to assess the role of PfPK2 in calcium release. For this purpose, parasites were loaded with cell permeable calcium-sensitive fluorophore Fluo-4-AM to measure free calcium in the parasite as described previously (Biagini et al, 2003; Nofal et al, 2021). Phosphodiesterase inhibitor zaprinast - which causes sustained calcium release from intracellular stores (Nofal et al, 2021; Paul et al, 2020), was used to treat PfPK2-loxP parasites. One set of parasites was also treated with calcium ionophore A23187 to estimate the amount of total stored calcium. RAP- treatment of PfPK2-loxP parasites caused a significant decrease in calcium release as indicated by the data from four independent biological replicates and almost no change in A23187 induced calcium release was observed. These data suggested that PfPK2 is involved in the release of calcium from intraparasitic stores (Fig 7D).

Collectively, present studies shed light on a novel signalling pathway in malaria parasite in which PfPK2 is an upstream regulator of key events like cGMP synthesis, which in turn is critical for calcium release. Given these second messengers are critical for host erythrocyte invasion, it is reasonable to conclude that PfPK2 regulates invasion by controlling the levels of these second messengers in the parasite, which in turn activate downstream events involved in key processes like microneme secretion (Fig. 8). In addition, PfPK2 may also regulate the glideosome-actomyosin motor activity as its depletion perturbs phosphorylation of key proteins involved in this activity (Fig. 6B and 6C, Table 1), although it needs further experimental validation.

**Figure 8.**
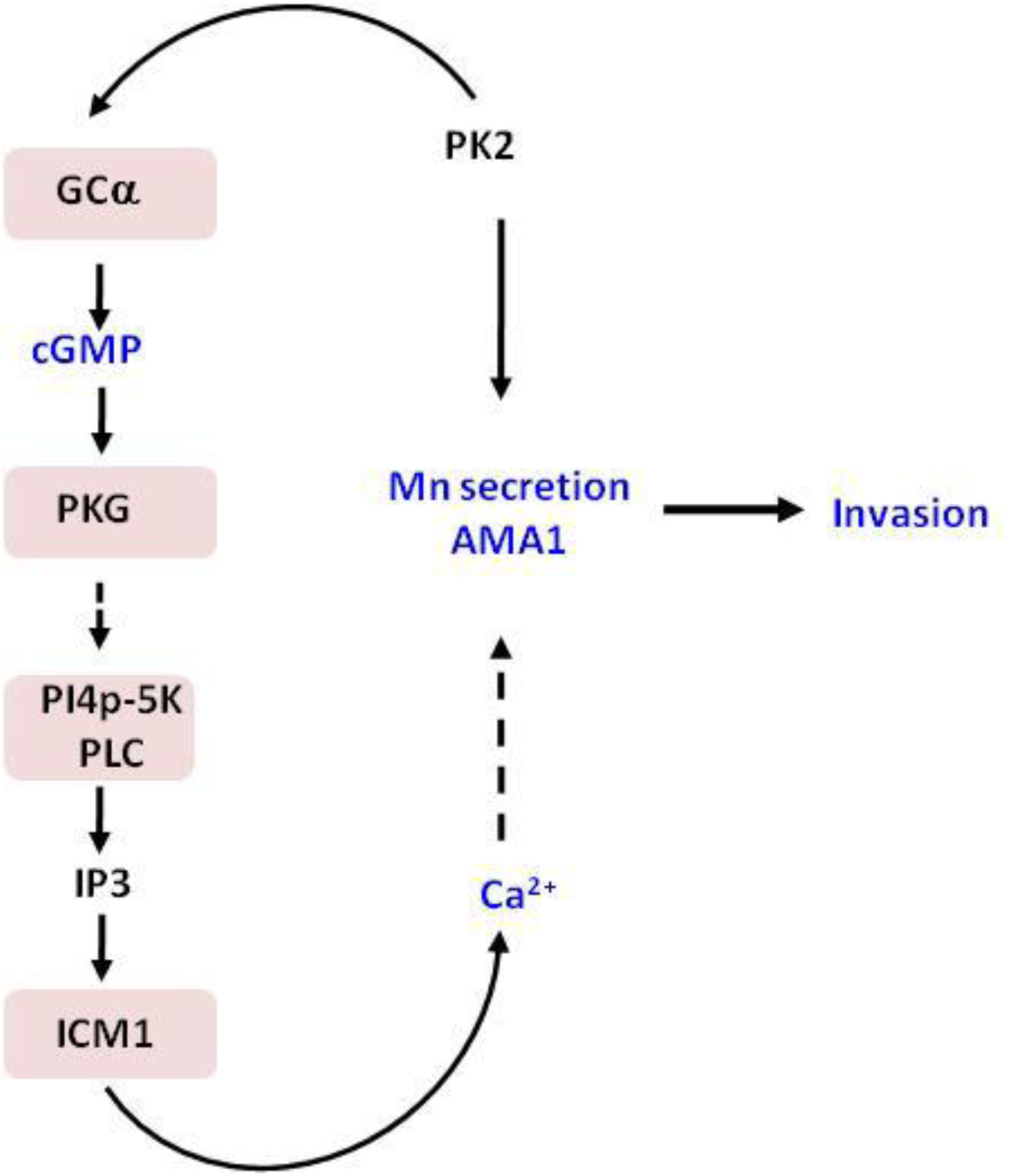
A novel PfPK2- dependent signalling pathway was identified in which PfPK2 may promote cGMP production (Fig. 7C) by regulating GCα (Fig. 6B, 7A), which in turn may facilitate the release of calcium (Fig. 7D) from intraparasitic stores and activate PfPKG, which is known to trigger calcium release in the parasite (Baker et al, 2017a; Brochet et al, 2014). Calcium, which is critical for the invasion, may also facilitate the release of microneme proteins like AMA1 (Fig. 4), which is important for invasion as it regulates the Tight Junction between the parasite and the host erythrocyte.

## Discussion

Present studies demonstrate that PfPK2 is critical for host erythrocyte invasion by *P. falciparum*. PfPK2 seems to be present in compartments that are proximal to the micronemes and rhoptries but did not exhibit co-localization with proteins that reside in this organelle. The compartment in which it is present appears to be vesicular as indicated by its punctate staining and each merozoite seems to contain 2-3 PfPK2-positive puncta, which do not co-localize with apical organelles. It is possible that its localization to these vesicular structures is dynamic and it is trafficked to other locations transiently, which may allow it to target proteins at distinct locations. PfPK2 depletion resulted in a severe impairment in microneme secretion, which included the release of AMA-1. Its depletion also caused defects in secretion of other microneme protein EBA175, which suggested that PfPK2 may have a generic role in microneme release. The parasite utilizes multiple pathways for the invasion of host erythrocytes, it interacts with the sialic acid on glycophorin A via EBL family members like EBA-175 that are released from the micronemes (Duraisingh et al, 2003; Sim et al, 1994). Alternatively, it utilizes sialic acid independent pathways, which typically involve RH family members like RH5 (Crosnier et al, 2011). Despite reduced EBA175 secretion, the attachment was almost unaltered in PfPK2-depleted parasites, which suggested that independent pathways like RH5-basiginin may operate effectively to facilitate its attachment.

In contrast, the entry of the parasite into the erythrocyte was significantly compromised and corroborated well with reduced AMA-1 secretion. Live cell imaging revealed that PfPK2- depleted parasites exhibited prolonged attachment to erythrocytes but failed to invade or enter the erythrocytes. However, characteristic deformation of the erythrocyte membrane and/or echinocytosis was observed. These defects share striking resemblance to observations made in studies in which AMA1 was depleted or parasites were treated with R1 peptide, which prevents interaction between AMA1 and RON2/4 (Richard et al; Treeck et al, 2009; Yap et al, 2014). IFA studies revealed that PfPK2 depleted merozoites that were attached to the erythrocyte surface exhibited either very low levels or absence of detectable AMA1 at the TJ whereas RON2 was unchanged (Fig. 5C), which supported a possible role of PfPK2 in TJ formation.

It was interesting to note that a few PfPK2-loxP merozoites managed to invade the erythrocytes upon RAP-treatment. However, most of these rings either did not mature and/or turned pyknotic (Fig. 2F and 2G). It is indeed possible that post invasion, sealing of erythrocyte membrane may be impaired due to issues like abrogated TJ formation etc. As a result, the environment for the newly formed rings inside the host erythrocyte was possibly not congenial for growth.

In order to understand the mechanism via which PfPK2 may regulate host cell invasion, comparative phosphoproteomics was performed to identify proteins which may be regulated by PfPK2 signalling. These studies resulted in identification of several putative targets which may be involved in processes that may be pertinent to PfPK2 function in invasion which included signalling, invasion and glideosome/actomyosin motor (Fig. 6B and 6C, Table 1, Supp Data Set S1). Previously, the role of PfGAP45 in host cell invasion has been demonstrated (Perrin et al, 2018) and its phosphorylation may promote its function of actomyosin motor anchoring (Thomas et al, 2012). Several other proteins of the motor and glideosome complex like Myosin E, Myosin light chain, GAP40 and IMC proteins also exhibited altered phosphorylation in PfPK2-depleted parasites suggesting that it may directly or indirectly facilitate their phosphorylation. The glideosome complex facilitated motor activity is also be required for movement of the merozoite post-TJ formation from apical to the basal end (Yahata et al, 2021). It is possible that altered phosphorylation of PfGAP45 and other glideosome-actomyosin related targets may contribute to this process.

One of the features of comparative phosphoproteomic studies was the fact that several signalling proteins exhibited aberrant phosphorylation upon PfPK2 depletion: Adenylyl cyclase β and PfPKAc were hyper-phosphorylated, which is likely to be an indirect effect possibly caused by deregulation of a phosphatase. Incidentally PPM2 phosphorylation was also modulated. Guanylyl cyclase (GCα)-which generates cGMP in the parasite especially in the blood stages- is involved in several important processes like egress and invasion via its downstream effector kinase PfPKG (Nofal et al, 2021). Since GCα phosphorylation at S1214 was attenuated upon PfPK2 depletion, cGMP formation was evaluated, which confirmed the involvement of PfPK2 in cGMP generation. As mentioned above, this residue is unique to GCα from *Plasmodium spp.* as it is in an insert of the N-domain (Fig. 7A). Since this domain is important for ATP binding (Nofal et al, 2021), phosphorylation of this insert may have a bearing on the function of ATPase domain. Interestingly, GCα was previously reported to be hyperphosphorylated at S674 upon depletion of protein phosphatase PfPP1 and cGMP levels were reduced upon depletion of this phosphatase (Paul et al, 2020). These phosphorylation events also occur in the ATPase domain, which was reported to play a role in cGMP synthesis and function of PfGCα (Nofal et al, 2021). While the direct relevance of phosphorylation of these sites in GCα function remains to be established, these studies suggested that its phosphorylation may play a critical role in its function of cGMP generation. One of the major functions of cGMP-PfPKG signalling is to stimulate the release of calcium in the parasite, which in turn regulates these processes (Bagur & Hajnoczky, 2017; Brochet & Billker, 2016; Brochet et al, 2014; Clapham, 2007; Nofal et al, 2021). Indeed, PfPK2 depletion impaired cGMP levels in the parasite (Fig. 7C). Furthermore, calcium release was also significantly reduced in PfPK2-depleted parasites, which corroborated well with reduced cGMP levels (Fig. 7D). Given that calcium signalling is involved in invasion (Brochet & Billker, 2016; Kumar et al, 2017; Vaid et al, 2008b), it is reasonable to propose that one of the ways PfPK2 may regulate invasion may be by facilitating calcium release possibly by regulating cGMP signalling. While cGMP regulation of calcium release-which occurs via PfPKG- has been well established, the underlying mechanism are not very clear.

PI(4,5)P2 is hydrolyzed by PLC to generate IP3 and DAG and IP3 regulates calcium release. GCβ and PKG have been shown to regulate PIP2 and PIP3 levels in *P. berghei* ookinetes, which was attributed to possible phosphorylation of PI4K and PI4P5K by PbPKG as their phosphorylation was compromised upon depletion of this kinase (Brochet et al, 2014). In present studies, PI4P-5K phosphorylation was also reduced in the phosphorylation in PfPK2- depleted parasites (Fig 6B, Table 1, Supp Data Set S1). Therefore, it is possible that PfPK2 mediated cGMP generation has a bearing on PIP2 formation and/or metabolism, which contributes to calcium release in the parasite.

Present studies established the role of PfPK2 in host erythrocyte invasion and specifically in its ability to regulate late events in invasion possibly via TJ formation. A novel pathway was deciphered in which it is an upstream regulator of cGMP-calcium signalling axis, which is critical for invasion (Fig. 8). PfPK2 can be an attractive drug target as its inhibition is likely to block a signalling cascade necessary for the parasite to establish infection in the human host.

## Materials and Methods

### Parasite cultures

*Plasmodium falciparum* strains 1G5DC was obtained from European Malaria Reagent Repository (EMRR) and DSM1 was obtained from *BEI* resources, Malaria Research and Reference Reagents Resource Centre (MR4), American Type Culture Collection (ATCC). Parasites were maintained in O+ human erythrocytes (5% hematocrit) in RPMI-1640 supplemented with 0.5% Albumax II and 50 μg/mL hypoxanthine (Trager & Jensen, 1977). Cultures were maintained at 37°C under a mixture of 5% CO_2_, 3% O_2_, and 91.8% N_2_ or 5% CO_2_. Fresh erythrocytes were used to dilute parasite with fresh culture media to maintain 3-5% parasitemia at 5% haematocrit. For culturing various transgenic lines relevant drugs were used as described below. Parasite synchronization was carried out using sorbitol as described previously (Lambros & Vanderberg, 1979).

### Percoll purification

*P. falciparum* infected erythrocytes (containing ∼5% schizonts) were pelleted down and resuspended in 2 ml complete medium. 3 ml of 70% percoll was used and infected-erythrocytes were gently poured alongside the rim of the tube. The samples were centrifuged in a swinging bucket rotor for 10 min at 2000 rpm (RT) without brake. The parasites which were mainly enriched schizonts, were gently removed and washed once with cRPMI and added to a new flask containing fresh erythrocytes at 2% hematocrit.

### Plasmid construction and transfection into 1G5DC parasites

#### 1G5DC+pSLI-PfPK2-loxP

All PCR primers used in this study were synthesized by Sigma and are indicated Supplementary Table S1.

A transgenic parasite line using dimerazable Cre recombinase (diCre) was generated to knockdown PfPK2. These parasites were generated in 1G5DC background as these parasites express diCre. A selection linked integration (SLI) approach was employed for this purpose (Birnbaum et al, 2017). For this purpose, a 330bp homology arm corresponding to 5’-end of *PfPK2* gene was amplified using primers 1/2 and cloned in *NotI* and *PmeI* sites upstream of loxP and Myc-tag coding sequence in pSLI-N-sandwich-loxP(K13) vector, which was a gift from Tobias Spielmann (Addgene plasmid #85793; http://n2t.net/addgene:85793; RRID:Addgene_85793). A recodonized version of full-length PfPK2 with at *AvrII* and *StuI* sites upstream of a loxP sequence in frame with GFP tag was custom synthesized as a G-block (Genscript). The resulting plasmid DNA construct (pSLI-PfPK2-loxP) was transfected in 1G5DC *P. falciparum* strain (Collins et al, 2013a). Transfections were performed using ∼100μg of purified plasmid DNA constructs in uninfected erythrocytes followed by the addition of percoll-enriched schizont stage parasites. Subsequently, transgenic parasites were selected by using 2nM WR99210 and recombinants were enriched using 1.5µM DSM-1.

Drug selected parasites were subjected to genotyping followed to assess the modification of the desired locus. After limited dilution cloning of drug resistant parasites, a clone was obtained, which lacked the endogenous unmodified locus was used for further analysis. PCR for genotyping of was performed followed by Sanger sequencing for the PCR products to confirm the desired modifications. Rapamycin was added to dimerize the Di-Cre recombinase in 1G5DC/PfPK2-loxP transgenic line resulting in the excision of loxP flanked region in the modified PfPK2 genomic locus.

#### HA-cPfPK2-PK2-loxP

In the above-mentioned PfPK2-loxP parasites, PfPK2 was overexpressed to complement the effects of PfPK2 depletion. For this purpose, PfPK2 cDNA was prepared from *P. falciparum* 3D7 RNA (isolated using QIAGEN RNeasy kit) and PCR amplification was done using primers 10/11 to amplify PfPK2 with an N-terminal HA tag and primers 8/9 was used to mutate the internal *KpnI* site in PfPK2 by overlapping PCR. The PCR product was cloned in pARL-BSD vector (a kind gift from Dr. Asif Mohmmed) using *KpnI* and *AvrII* restriction sites. The plasmid DNA was transfected as described below, followed by selection using 2.5μg/ml of blasticidin and 1.5 μM DSM1.

### Growth rate and invasion assays

PfPK2-loxP parasites were cultured in the presence of 5 nM WR99210 and 1.5 μM DSM1. Subsequently, for growth rate assay, ring stage synchronized parasites were seeded at ∼0.5-1% parasitemia at 2% hematocrit. 200 nM rapamycin was added to parasite cultures for ∼6-8 hours to deplete PfPK2. Parasite growth was monitored periodically by microscopic analysis of Giemsa-stained thin blood smears and counting individual parasitic stages.

For flow cytometry based assays, samples were collected at desired time and fixed with 1% PFA/0.0075% glutaraldehyde and kept on an end-to-end rocker for 15 min. After completion, samples were either stored at 4°C or processed directly for Hoechst 33342 staining for 10 min at 37°C. Samples were then washed at least twice with FACS buffer (0.3% BSA + 0.02% Sodium Azide) followed by analysis (Bei et al, 2010; Theron et al, 2010) on BDverse (BD biosciences).

The parasite invasion assays were carried out as described previously with slight modifications (Ekka et al, 2020; Kumar et al, 2017; Theron et al, 2010). Schizonts were washed twice and resuspended in culture medium with fresh erythrocytes to achieve 2% hematocrit. Invasion experiments were performed using 0.5-1% schizont parasitemia and 2% hematocrit in a 6-well plate in a gas chamber equilibrated with gas mixture described above at 37°C and samples were collected periodically for upto ∼12 hours. Thin blood smears were Giemsa-stained and the number of schizonts or newly formed rings was counted. Alternatively, flow cytometry was used as described previously (Ekka et al, 2020; Theron et al, 2010) for which parasites were fixed using 1% PFA + 0.0075% glutaraldehyde in either ALS solution or in FACS buffer followed by staining the parasite nuclei using Hoechst 33342. Flow cytometry data was acquired using BD FACS verse and analysed using FlowJo V.10 software (Tree Star, Ashland, OR).

### Live-cell Imaging

Time-lapse imaging was performed as described previously with slight modifications (Ekka et al, 2020). Briefly, PfPK2-loxP parasites were synchronized at the ring stage and treated with DMSO or rapamycin for 6h and were allowed to develop to schizont stage. Schizonts purified and were cultured in the presence of 25nM PfPKG-inhibitor ML10 (Baker et al, 2017b; Ressurreicao et al, 2020) with fresh erythrocytes at 2% hematocrit in RPMI1640 complete media. Subsequently, ML10 was washed after 4-6h and parasites were plated on glass bottom chamber slides (Ibidi) coated with 0.5 mg/ml concanavalin A. The slides were kept at 37°C, supplied with a humidified environment under 5% CO_2_ and imaging was performed using a Zeiss Axio Observer microscope. Images were typically taken every 2s for at least 10 min, and resulting time-lapse videos were processed using Axio Vision 4.8.2 analysis software.

### Parasite attachment assay

The parasite-erythrocyte attachment assays were carried out as previously described with minor changes (Kumar et al, 2017; Paul et al, 2015). PfPK2-loxP parasites were synchronized at ring stage (∼3% parasitemia, 2% hematocrit) and treated with DMSO and Rapamycin as described above and were allowed to mature to schizont stage (∼44 hpi). Subsequently, 25nM ML10, which prevents egress was added for 6h, to allow maturation of schizonts and merozoites were released subsequent to the washing of the inhibitor (Ressurreicao et al, 2020). Assays were performed in 6-well culture plates containing fresh erythrocytes treated either with 1 mM cytochalasin D or DMSO (control) with shaking at 80 rpm at 37°C. After 8-12 h, thin blood smears were stained with Giemsa. The number of parasites attached to erythrocytes was quantified by microscopically examining the stained parasites. In some experiments, IFA was performed to visualize the tight junction as described below.

### Phosphoproteomics and Mass pectrometry

#### Proteomics Studies

##### Cell lysis and protein extraction and TMT labeling

The 1G5DC or PfPK2-loxP parasites were cultured and synchronized using sorbitol. Ring stage parasites were treated with either DMSO or RAP for duration of 6 hours. The parasites were subsequently harvested upon maturation to schizonts, which was done ∼44 hours post-infection, by lysing the infected red blood cells with saponin. The resulting parasite pellet was washed and dissolved in protein extraction buffer (50 mM triethylammonium bicarbonate (TEABC), 2M SDS, and 1X protease and phosphatase inhibitors). The samples were sonicated and centrifuged, and the resulting supernatant was used for protein estimation by bicinchoninic acid assay. Equal amounts of protein were taken from both DMSO and RAP-treated parasites and subjected to reduction with 20 mM dithiothreitol (DTT) at 60°C for 30 minutes, followed by alkylation with 20 mM iodoacetamide for 10 minutes in dark at ambient temperature. Subsequently, proteins were digested using a trypsin and lys-C mix (Promega, Madison, USA) at a ratio of 100:1, and tryptic peptides were cleaned using a C18 column and dried with a SpeedVac concentrator (Thermo Fisher Scientific, USA). The peptides were quantified using the Pierce Quantitative Colorimetric Peptide Assay Kit. Subsequently, 500 µg of peptides from each sample were taken and processed for TMT labelling as per manufacturer’s instructions. After confirming a labelling level of >95%, the reaction was quenched by adding 8 µl of hydroxylamine. The labelled peptides were pooled, dried, and cleaned using C18 Sep-Pak (Waters, Milford, MA, USA).

##### Phosphopeptide enrichment and fractionation

A 50µg aliquot of the sample was used for total proteome analysis, and the remaining sample was subjected to phosphopeptide enrichment. For the first phosphopeptide enrichment step, we used the TiO2 Phosphopeptide Enrichment Kit (Thermo Scientific, Catalog number: 88303), following manufacturer’s instructions. The flow-through collected during this step was used for the second phosphopeptide enrichment step, utilizing the High-Select Fe-NTA Phosphopeptide Enrichment (Catalog number: A32992). The eluent from both phosphopeptide enrichment steps was combined for fractionation. To resolve complexity, we employed C18 stage tip fractionation, collecting a total of 24 fractions that were subsequently concatenated into six fractions. The peptide samples were dried and dissolved in 0.1% formic acid prior to mass spectrometry analysis.

##### Mass spectrometric analysis

Mass spectrometry analysis was performed using an Orbitrap Fusion Tribrid mass spectrometer coupled to an Easy-nLC 1200 nano-flow UPLC system (Thermo Fischer Scientific, Germany). All fractions were loaded onto a trap column nanoViper (2 cm, 3 µm C18 Aq) (Thermo Fischer Scientific). Subsequently, eluted peptides were further resolved on an analytical column (75 µm × 15 cm, C18, 2 µm particle size) (Thermo Fischer Scientific). A gradient of 5-35% solvent B (80% acetonitrile in 0.1% formic acid) was used over a 120- minute, at a flow rate of 300 nl/min.

Data were acquired in data-dependent acquisition (DDA) mode. Precursor ions were acquired in full MS scan mode in the range of 400-1600 m/z, with an Orbitrap mass analyzer resolution of 120,000 at 200 m/z. An automatic gain control (AGC) target value of 2 million with an injection time of 50 ms and dynamic exclusion of 30 seconds was used. The selection of the most intense precursor ions was performed at top speed data-dependent mode, isolated using a quadrupole with an isolation window of 2 m/z and an isolation offset of 0.5 m/z. Subsequently, filtered precursor ions were fragmented using higher-energy collision-induced dissociation (HCD) with 34±3% normalized collision energy. MS/MS scans were acquired in the range of 110-2000 m/z in the Orbitrap mass analyzer at a mass resolution of 60,000 mass resolution/200 m/z. AGC target was set to 100,000 with an injection time of 54 ms. For each condition, MS/MS data were acquired in duplicates.

##### Data analysis

MS/MS raw data was used to search against a combined protein database of *P. falciparum* 3D7 (downloaded from PlasmoDB web resource, version 46) and humans using SEQUEST and Mascot (Version 2.4.1) via Proteome Discoverer v2.2 (Thermo Fisher Scientific). Trypsin was selected as the protease of choice with a maximum of two missed cleavages. A precursor mass tolerance and fragment mass tolerance of 10 ppm and 0.02 Da were selected, respectively and the data was filtered at 1% FDR at PSM level. Static modifications were carbamidomethylation at cysteine and TMT labeling at lysine and N-terminus of peptide. For dynamic modifications, methionine oxidation, acetylation at N-terminus of protein and S/T/Y phosphorylation were selected. The phosphorylation fold changes were normalized against the total proteome data as mentioned earlier (Bansal et al, 2021). The ptmRS node was used during the search to determine the probability of phosphorylation site localization and a ptmRS score ≥99% was used as a cut off.

Phosphopeptides along with their fold changes (RAP/DMSO) from three biological replicates were uploaded on Perseus software (Version 1.4) to determine the p-value using student’s t-test for each phosphosite and p<0.05 was considered as significant (Supplementary Data Set S1 for PfPK2-loxP parasites and Data Set S2 for 1G5DC control line). In addition, fold change cut-off of 1.2 to further filter differentially phosphorylated sites (Kumar et al, 2017). The prediction of protein-protein interactions was performed using, STRING web resource (Version 11.5) was used. Proteins that were found to be differentially phosphorylated were considered for the analysis, and interactions were predicted with medium confidence filter (Szklarczyk et al, 2011).

### Immunofluorescence Assay (IFAs)

Immunofluorescence assays (IFA) were performed either on parasite suspensions or thin blood smears as previously described (Tonkin et al, 2004). For IFA on cell suspension, the parasite pellet was washed in 1x PBS to remove the culture medium and fixed for 30 min using 4% PFA and 0.0075% glutaraldehyde prepared in PBS. For permeabilization, 0.1% Triton-X100 was used for 15 min. For air-dried thin blood smears, cold methanol and acetone mix (1:1, v:v) was used for 2 min followed by blocking with 3% BSA for 45 min at room temperature was used for blocking followed by incubation with primary antibody for 12h at 4 ^ͦ^C. After washing, Alexafluor mouse/rabbit 488/594-labeled secondary antibodies were added (Invitrogen) for 2h at ambient temperature. Samples were mounted using Vectashield mounting media (Vector Laboratories Inc.) which contained DAPI to label nuclei. The stained parasites were visualized using either Axio Imager Z1 microscope or LSM980 confocal microscope (Carl Zeiss). The processing of images was done using AxioVision 4.8.2 or Zeiss ZEN black/blue software. Z-stacks that best represented the immunolocalization were used for illustrations in the figures.

### Immunoblotting

Parasites were collected and infected erythrocytes were lysed using 0.05% saponin (w/v) on ice for 10 min. Centrifugation at 8000 rpm was done to isolate the parasite pellet, which was washed three times with pre-chilled PBS. The parasite pellet was re-suspended in lysis buffer (10 mM Tris pH 7.5, 100 mM NaCl, 5 mM EDTA, 1% Triton X-100, and complete protease inhibitor cocktail (Roche Applied Science) or 2% SDS and homogenized by either passing the solution through a 26-gauge needle or by sonication. The supernatant from the centrifugation of the lysates at 14,000 g for 30 min at 4°C was used to estimate the amount of protein using a BCA protein estimation kit (Pierce). After separation by SDS-PAGE, lysate proteins were transferred to a nitrocellulose membrane. Immunoblotting was performed as described previously (Ekka et al, 2020; Kumar et al, 2017) using specific antisera and blots were developed using SuperSignal^®^ West Pico or Dura Chemiluminescence Substrate (Thermo Scientific) following manufacturer’s instructions.

### Detection of parasite secreted proteins

PfPK2-loxP parasites were synchronized tightly at ring stage and treated with DMSO and rapamycin followed and were allowed to mature to schizonts (42-44 hpi) and secrete microneme and rhoptry proteins were detected in the extracellular milieu as described previously (Ekka et al, 2020; Kumar et al, 2017). The residual supernatant was used for Western blot analysis with antibodies against several proteins of interest, as mentioned below.

### Expression of recombinant proteins

For the expression of recombinant proteins and subsequent kinase assays, *PfPK2* was amplified using primer 12/13 from a plasmid that contained custom synthesized codon optimized version of PfPK2. The amplicon was cloned in *BamHI* and *NotI* restriction sites of pET28a (+) vector. The deletion mutants of PfPK2 protein were generated either by site directed mutagenesis or overlapping PCR using primers indicated in Table S1. Briefly, to generate K140M and ΔRD mutant of PfPK2 site directed mutagenesis was performed using primer 14/15 and 16/17, respectively. PfPK2 deletion mutant for ΔC and ΔRD+ΔC was generated using primer set 12/18 and 12/19, respectively. The final PCR product was cloned in pET28a vector under *BamHI* and *NotI* restriction site as mentioned above.

Recombinant proteins were expressed in BL-21 (DE3) RIL strain using 1 mM of IPTG at 18°C for 16 h. Subsequently, the bacterial culture was pelleted at 3500 rpm for 15 min at 4°C. Cells were resuspended in a solution containing 50mM potassium phosphate buffer pH 7.4, 200mM NaCl, 1% Triton X-100, 10% glycerol, 1mM β-mercaptoethanol, and protease inhibitors. Sonication was performed for 10 min on ice followed by centrifugation at 10,000 rpm for 45 min to separate the supernatant and pellet fraction. The supernatant was collected and used for recombinant protein purification either by using AKTA FPLC or on His-Trap columns (GE Healthcare) as per the manufacturer guidelines or by manually purifying pre-equilibrated Ni-NTA-agarose (Qiagen). Purified proteins were eluted with imadzole and dialyzed overnight at 4°C against 50 mM Tris, pH 7.4, 10% glycerol, and 1 mM DTT.

### Kinase assays

Tyically, protein kinase assays were performed using recombinant ∼0.1 μM 6xHis-PfPK2 in a buffer containing 50 mM Tris pH 7.5, 10 mM magnesium chloride, 1 mM DTT and 100 µM [γ-^32^P] ATP (6000 Ci/mmol) (Perkin Elmer). In some experiments, 5 μM calmodulin in the presence of 10 μM CaCl_2_ or 100 μM EGTA was used and 2 μg of Histone IIa was used as phosphoacceptor substrate. Typically, the reactions were carried out at 30° C for 45 min in a water bath and were stopped by boiling the reaction mix in SDS-PAGE sample buffer. The samples were subjected to SDS-PAGE, the gel was subsequently dried and exposed to a phosphorimager image plate. The phosphorylation of the substrate was then detected by phosphorimaging using an Amersham Typhoon scanner (GE Healthcare). Equal amount of PfPK2 and various mutants were used to compare their activity in assays which were performed as described above.

### Determination of calcium release in the parasite

Calcium release in the parasite was measured as described previously with brief modifications (Nofal et al, 2021). ∼1×10^7^ percoll purified schizonts from DMSO- or RAP-treated cultures were incubated in a phenol red-free RPMI containing 10μM Fluo4-AM (Invitrogen, Carlsbad, CA) in dark at 37°C for 45 min. Parasites were washed with phenol red-free RPMI and after 20 min 100μl of parasite suspension was transferred to a 96-well plate. As a control, in some wells only phenol red-free RPMI was included. The fluorescence was measured at 22-sec intervals for 5 min using a Synergy H1 microplate reader (BioTek) with excitation and emission wavelengths of 483 and 525 nm, respectively. Parasites were resuspended and transferred to wells containing 1.5% DMSO or 75μM zaprinast or 20μM A23187 (20μM) and fluorescence was measured as described above. The average of the relative fluorescence units at each time point and condition was determined after subtracting baseline and DMSO control values.

### Measurement of intracellular cyclic GMP levels

Intracellular cyclic nucleotide levels in mature segmented schizonts were measured using enzyme-linked immunosorbent assay (ELISA)-based high-sensitivity cGMP assay kit as suggested by the manufacturer (Cell Signalling Technology, Cat No. #4360). Briefly, ∼10^7^ percoll-purified schizonts were obtained from RAP- or DMSO-treated cultures. The purified schizonts were incubated for 30 min in culture medium containing ML10 in the presence of zaprinast (100 μM) (Nofal et al, 2021). Parasites were pelleted at 9,000 × g, washed with 1x PBS followed by centrifugation at 9,000 × g, and the pellet was collected and stored at –80°C until required. Parasites were lysed using cell lysis buffer, as per the manufacturer’s instruction and standards were also prepared in the same buffer. 50 μl samples and 50 μl HRP-linked cGMP solution was added to the cGMP assay plate and incubated for 3h on horizontal orbital plate shaker at room temperature followed by 4 washes with 200 μl wash buffer. Further incubations were performed in 100 μl of TMB substrate for 30 min at ambient temperature. Reaction was terminated by adding 100 μl STOP solution and absorbance was measured at 450 nm using a Synergy H1 microplate reader (BioTek).

### Densitometry and statistical analysis

Image J (NIH) software was used to perform densitometry of Western blots. The band intensity of the loading control was used for normalization. Statistical analysis was performed using Prism (Graph Pad software Inc USA). Data was generally represented as mean ± Standard error of mean (SEM), unless indicated otherwise and p < 0.05 was considered as statistically significant.

## Data availability

The mass spectrometry-based proteomics data have been deposited to the ProteomeXchange Consortium (http://proteomecentral.proteomexchange.org) via the PRIDE partner repository with the dataset identifier PXD042264. The data can be accessed using following details: Username: Password: rUQtE6o8

## Acknowledgements

Studies were supported by grants BT/COE/34/SP15138/2015 from Department of Biotechnology, a Team Science grant (IA/TSG/21/1/600261) to PS and TSKP from DBT-Wellcome Trust India Alliance and funds from NII core. P.S. is a recipient of J.C. Bose Fellowship. The efforts of Puranjaya Pancholi and Prabneet Kaur in the expression of some of the recombinant proteins are appreciated.

